# PANoptosis, a combination of inflammatory cell death mechanisms, induced by Ophiobolin A in breast cancer cell lines

**DOI:** 10.1101/2025.04.08.647841

**Authors:** Santhalakshmi Ranganathan, Tolulope Ojo, Alagu Subramanian, Jenna Tobin, Kayla L. Haberman, Alexander Kornienko, Angela Boari, Antonio Evidente, Mary Lauren Benton, Daniel Romo, Joseph H. Taube

## Abstract

An unmet challenge in managing breast cancer is treatment failure due to resistance to apoptosis-inducing chemotherapies. Thus, it is important to identify novel non-apoptotic therapeutic agents. Several non-apoptotic programmed cell death pathways utilize specific cellular signaling events to trigger lytic and pro-inflammatory cell death. PANoptosis, which encompasses pyroptosis, apoptosis and necroptosis, is of paramount importance in the regulation of cell death and immune responses. Our study illustrates that ophiobolin A (OpA) is an anti-cancer agent that triggers lytic cell death in breast cancer cells, including triple-negative breast cancer (TNBC), via a mechanism dependent on RIPK1. This study reveals that OpA induces typical pyroptosis-like characteristics, including cellular swelling, plasma membrane rupture, GSDMD cleavage and release of cytokines in breast cancer cells. The involvement of caspase 3, RIPK1, and GSDMD suggests that PANoptosis is activated upon OpA treatment in breast cancer. The induction of pro-inflammatory cell death suggests potential applications for OpA in cancer treatment.

## Introduction

Breast cancer is now the most prevalent form of malignant cancer, surpassing lung cancer, with approximately 2.3 million new cases diagnosed annually (1). Despite considerable advancements in adjuvant therapy for breast cancer treatment, the widespread resistance to conventional pro-apoptotic treatments presents a significant challenge (2, 3). Resistance is associated with an elevated risk of recurrence and metastasis, resulting in high mortality rates (4). There is a pressing demand for novel approaches to induce non-apoptotic cell death pathways to meet this clinical need.

Programmed cell death (PCD) is an essential event required in normal development and tissue homeostasis. The mechanism of PCD invoked by distinct drugs includes non-lytic (apoptosis, ferroptosis, paraptosis), and lytic forms (pyroptosis, necroptosis), and is highly relevant to anti-cancer treatment (5). Apoptosis proceeds through caspase (Casp) cleavage, specifically Casp3, Casp6, Casp7 and is initiated by Casp8 (6, 7). Ferroptosis has recently been identified as an essential type of cell death characterized by iron overload and lipid-peroxidation (8). Paraptosis is characterized by cytoplasmic vacuolation with dilation of the endoplasmic reticulum (9) and is distinct from apoptosis due to the absence of nuclear fragmentation, chromatin condensation, and caspase-dependency. Pyroptosis is dependent on caspase-1 activation, and exhibits cell swelling, membrane rupture, and the release of pro-inflammatory cytokines (10). Pyroptosis is also characterized by chromatin condensation and DNA fragmentation, although it differs morphologically from apoptosis by having an intact nucleus (11). Casp3, the protein responsible for executing apoptosis, can cause the plasma membrane to undergo secondary necrosis or pyroptosis by inducing membrane pore formation (12). Gasdermin E (GSDME) has also been demonstrated to convert TNF-α or chemotherapy drug-induced casp3-mediated apoptosis to pyroptosis (13). Given the complex features and interconnected signaling networks among these mechanisms it is perhaps not surprising that features of apoptosis, necroptosis, and pyroptosis, termed PANoptosis, may be coordinately activated (14). In the context of breast cancer, the potential to leverage pyroptosis/PANoptosis as a mechanism of cell death for therapy is promising, particularly in the context of resistance to apoptosis and for the capacity to synergize with immunotherapy approaches.

Ophiobolin A (OpA) is a sesterterpenoid produced by the pathogenic fungi of the *Bipolaris, Aspergillus, and Drechslera* genera that displays potent antiproliferative activity against multiple cancer cell lines, including chronic lymphocytic leukemia, breast cancer stem cells, and apoptosis-resistant glioblastoma (15-18). However, both the mechanism of action and type of cell death pathway invoked by OpA have been difficult to establish and are likely cell-type dependent. OpA has been shown to induce paraptosis-like cell death, a form of non-apoptotic cell death, in human glioblastoma cells (19), necrosis in osteosarcoma cells (20) and autophagy in human melanoma cells (21). We recently demonstrated that the TNBC cell line MDA-MD-231 is significantly more sensitive to the cytotoxic effect of OpA than the ER-positive breast cancer cell line MCF7 (22); however, the mechanism of cell death which may underlie this difference is not established.

Breast cancer is a grouping of molecularly distinct diseases with distinct cells of origin and molecular markers. While patients with HER2-enriched and hormone-receptor (HR)-positive breast cancers benefit from targeted treatments, patients with triple-negative breast cancer (TNBC) face more limited treatment options. In this study, we investigated the mechanisms of PCD induced by OpA treatment in multiple breast cancer cell lines representing specific molecular subtypes of the disease. Our study reveals that the mechanism of programmed cell death elicited by OpA is highly cell type dependent. In HER2-positive breast cancer cells OpA induces pyroptosis, while TNBC cell lines exhibit variable features of cell death pathways. These results illuminate the capacity of OpA to activate non-apoptotic PCD, particularly PANoptosis, against breast cancer cell lines. This is the first study showing Ophiobolin A to induce a combination of non-apoptotic PCD in breast cancer.

## Materials and methods

### Cell culture and stable cell line

Human breast cancer cell lines MCF7, T47D, MDA-MB-231, MDA-MB-468, Hs578T, SKBR3 as well as non-transformed mammary cell line MCF10A, and fibroblast cell line 293T were purchased and cultured as recommended by American Type Culture Collection (ATCC). All the cell lines were cultured with respective media (Gibco, USA) contained 10% Fetal Bovine Serum (FBS, Gibco, USA) and 1% penicillin and streptomycin in a humidified incubator at 37°C and 5% CO_2_. All cell lines were routinely tested for the absence of mycoplasma contamination. HeLa cells and plasmid expressing RIPK3 were a kind gift of Zhigao Wang (23).

### Ophiobolin A source and purification

OpA was produced by *Drechslera gigante*a and purified from the fungal culture filtrates as reported previously (24). Briefly, the fungus was grown and maintained on Petri dishes containing PDA (Oxoid, England). The culture filtrates were lyophilized, redissolved in distilled water and extracted with EtOAc. The combined organic extracts were dehydrated, filtered, and evaporated under reduced pressure. The brown oily residue was fractionated by column chromatography and crude OpA was crystallized as white needles.

### Cell culture treatments and viability assay

To assess the cell viability, cells were seeded at the density of 5000 cells per well in 96-well plates. Compounds used include: zVAD (Selleck chem Cat#S7023), Ferrostatin-1 (Sigma-Aldrich Cat#SML-0583), Spautin-1 (MilliporeSigma Cat#56756910MG), Disulfiram (MCE Cat#HY-B024), Necroststin-1 (ThermoScientifric Cat# J65341FPL), GSK872 (Selleckchem), GSK840 (Selleckchem), and GSK843 (Selleckchem). Cells were treated with OpA in presence or absence of the inhibitors for 24h, later Cell Titer Blue (CTB) (Promega G8081) was added directly to each well according to manufacturer protocol, incubated for 3h at 37°C and fluorescent signal was recorded at 560_Ex_/590_Em_. Cells treated with combination of TNF-α (MilliporeSigma Cat# GF023), Smac/LCL161 (MCE CAS#1005342-46-0) and zVAD were used as positive controls for necroptosis.

### Autophagy assay

Cells were cultured to 85% confluence, dosed, washed with PBS and stained with CYTO-ID Green Detection Reagent according to manufacturer’s instructions (Enzo Life Sciences Cat# 1750200). The individual cellular fluorescence was measured using the Lionheart FX Automated Microscope (BioTek) under the cellular analysis protocol. The fluorescence captured was then quantified and analyzed using GraphPad Prism.

### Annexin V assay

For evaluation of apoptosis, cells were labelled using Annexin V-FITC early apoptosis detection kit (Biolegend Cat#640914) following the manufacturer’s protocol. Briefly, cells were plated on 10 cm dishes and treated with or without OpA or staurosporine (1µM) for 24h. Following trypsinization, the harvested cells were pelleted and resuspended in Annexin V FITC in binding buffer and incubated for 30 min at room temperature in dark and then propidium bromide (PI) was added for FACS analysis using BeckmanFACS Melody (BD Biosciences).

### Live cell imaging

To interrogate the formation of necrosomes, RIPK3 overexpressing cell were seeded in 96 well glass bottom plates (Corning Ref4580), treated with DMSO or OpA or TNF-α (Thermo Scientific Cat#YD370435) and zVAD along with doxycycline (FisherBioReagents Cat# BP26535) for 24h. To examine the integrity of the plasma membrane, the cells were treated with or without OpA for 4h and stained with Cellbrite550 (Biotium Cat#30105A) and SYTOX green (Invitrogen Cat#S7020) in respective media for 30 min at 37 °C. Later the cells were imaged using confocal microscope. Cell treated with Thapsigargin (EMD Millipore Cat#586005) were used as positive controls for necroptosis.

### Transmission electron microscopy (TEM)

Cells were grown to 80% confluency as described above and exposed to OpA for 4h. The cells were fixed with 2.5% glutaraldehyde dissolved in 0.06 M phosphate buffer (pH 7.2) for 30 min. The samples were washed thrice with PBS for 10 min, followed by a secondary chemical fixation for 30 minutes with 1% osmium tetroxide and 0.8% potassium ferrocyanide buffered with PBS (pH 7.2). Reagents were aspirated and the dish was washed thrice with PBS for 10 min. The sample then underwent a serial dehydration in ethanol: one incubation of 50% EtOH for 10 min, one incubation of 70% EtOH for 10 min, two incubations of 90% EtOH for 10 min, and two incubations of 100% EtOH for 10 min. The sample was infiltrated with a mixture of epoxy resin (EMbed 812) and ethanol through three infiltration stages (1:2, 1:1, 2:1) and polymerized in 100% Embed 812 at 60°C for 48 hours. Following polymerization, samples were trimmed, sectioned to 80 nm, and placed on a copper grid. Post-staining was completed with 1% lead citrate for 5 min and 1% uranylic acetate for 15 minutes. The samples were imaged using transmission electron microscopy (ThermoFisher Spectra 300 S/TEM).

Endoplasmic reticula were measured manually on TEM micrographs with the freehand tool in Olympus CellSens Dimension software (Olympus America Inc., Version 2.2). Cells were labeled prior to analysis to avoid duplication and used once for analysis. To account for the random orientation, a minimum of 10 different and randomly chosen cells were measured per condition, with at least 200 endoplasmic reticula per condition. The data collected was exported for subsequent analysis by ANOVA using GraphPad v10.

### Immunoblotting

Cells were lysed with RIPA buffer (ThermoScientific Cat#J63306.AP) supplemented with 1X protease inhibitor (ThermoScientific Cat#78430) and 1X phosphatase inhibitor (ThermoScientific Cat#78428). The protein concentrations were determined using BCA reagent kit (ThermoScientific Cat#23225) and then equal amount (50µg) of whole cell lysates were subjected to 4-12% SDS-PAGE (GelScript Cat#M00653), transferred onto 0.45 µm PVDF membrane (ThermoScientific Cat#.88520). The membranes were incubated with appropriate primary antibodies for overnight. The primary antibodies used were as following: RIP3 (Cell Signaling 13526S), pRIP3 (Cell signaling Cat#93654S), MLKL (Proteintech Cat#66675-14g), pMLKL (Abcam Cat#ab187091), Caspase3 (Cell Signaling Cat#9662S), Gasdermin D (Abcam Cat#Ab215203), Actin (BD Bioscience Cat#612657), HRP-conjugated anti-mouse secondary antibody (Cell Signaling Cat#7076S) and anti-rabbit secondary antibody (Cell Signaling, Cat#7074S). Immunoblots were treated with ECL Prime (Cytiva) and imaged using Biorad ChemiDoc Imaging system. β-Actin was used as an internal standard. All primary antibodies were diluted 1:1000 with phosphate buffer saline tween (PBST) except actin (1:2000) and secondary antibody to 1:2000 dilution. Primary antibodies were incubated overnight at 4°C and secondary antibodies incubated for 2hr at room temperature.

### Reverse transcription and real time quantitative PCR

Total RNA was isolated from cells treated with or without OpA (1uM) for 6h, using TRIzol reagent (Ambion by Life Technologies #15596018) according to manufacturer’s protocol. cDNA was generated using 250 ng of RNA with the MultiScribe reverse transcriptase (ThermoFisher #4319983) according to the manufacturer protocol. The synthesized cDNA was used to perform real time quantitative PCR using qPCR blue mix (Azura genomics Cat#AZ2305) by Quantstudio5 system (Applied Biosciences). The comparative Ct method was used for relative mRNA quantification using the formula 2^-ΔΔCt^ (25) and beta-actin was used for normalization. All reactions were performed as at least three independent experiments. All primers were obtained from Integrated DNA Technologies (Newark, NJ, USA). Primers were added to a final concentration of 2.3 μM. Primer sequences were as follows: IL6, forward-TGAGGAGACTTGCCTGGTGA; reverse-CTGCACAGCTCTGGCTTGTT; human CXCL8/IL8, forward-ATGACTTCCAAGCTGGCCGT; reverse-TCCTTGGCAAAACTGCACCT, IL1B, forward-GCAAGGGCTTCAGGCAGGCCGCG, reverse-GGTCATTCTCCTGGAAGGTCTGTGGGC human ACTB, forward-CATGTACGTTGCTATCCAGGC, reverse-CTCCTTAATGTCACGCACGAT. All primer amplification undergone thermocycling condition of melting temperature 60°C. Samples were normalized to human β-actin (ACTB) as an endogenous reference. RT-qPCR was analyzed using the ΔΔC_T_ method (25). The data were analyzed in Microsoft Excel and Prism 6.0.

### Enzyme-linked Immunosorbent Assay (ELISA)

Conditioned media was collected from cell lines treated with or without OpA (1 μM) for 6h. The amount of human IL-8 protein in conditioned media was measured using human IL-8 ELISA kits according to the manufacturer’s protocol (RayBiotech Cat# ELH-IL8-1).

### Identification of differentially expressed genes

Library preparation and RNA-seq was conducted on triplicate samples using the services of Novogene. We used Salmon (26) to quantify the expression of transcripts from FASTQ files. We indexed the reads using the GRCh38 human genome with the following options: --threads 8 --gcBias –validate Mappings. Quantifications were imported to DESeq2 (v1.32.0) using the tximeta (v1.10.0) package in R (v4.1.1) (27, 28). DESeq2 was used to identify differentially expressed genes between the control and treated condition.

### Gene ontology and gene set enrichment analysis

Gene ontology analysis was performed using functional annotations for genes in the GRCh38 assembly of the human genome. We used the clusterProfiler (v4.0.5) package (29) to quantify overrepresentation of differentially expressed genes in all three Gene Ontology categories: Biological Process, Molecular Function, and Cellular Component. We also tested for enrichment of differentially expressed genes in KEGG pathways. We corrected for multiple testing using the Benjamini-Hochberg procedure and considered significance at a q-value < 0.05. REVIGO was used to generate a condensed set of GO terms by inputting GO terms with p-value < 10^−10^.

### Tumor growth

Female Scid/bg (CB17.Cg-PrkdcscidLystbg-J/Crl) mice (5–8 weeks old) weighing approximately 20g were obtained from Charles River Laboratories (Wilmington, MA, USA). Mice were maintained under a 12-hour light/dark cycle at a temperature of 20 °C to 22 °C. Food and water were available *ad libitum*. Mice were maintained in accordance with the Institutional Animal Care and Use Baylor University Committee procedures and guidelines. MDA-MB-231 cells were harvested, pelleted by centrifugation at 2000×g for 2 min, and resuspended in sterile serum-free medium supplemented with 30% to 50% Matrigel (BD Biosciences). Cells (2 × 10^6^ in 100 µl aliquots) were implanted into the left fourth mammary fat of 6–8-week-old mice and allowed to grow orthotopically until measurable by caliper. Then, vehicle or OpA (suspended in DMSO) or docetaxel (ThermoScientific AC456262500) was administered by intraperitoneal injection once a week for 6 weeks at 5 mg/kg OpA or 5 mg/kg docetaxel. Tumor volume and body weight were recorded concurrently with injection protocol. The progression of tumors was monitored every alternative days. Each group contained 4 to 6 mice. Humane endpoint for this study were established based on clinical observations including 20% decrease in body weight from the baseline, ulceration, tumor size beyond 2cm, visible signs of distress, dehydration or indication of any pain or discomfort in the animals. All the mice in the experiment were euthanized by CO_2_ inhalation and death was verified by lack of respiration and heartbeat.

### Statistical analyses

The experimental data were analyzed and plotted using GraphPad Prism 6.0. The mean between two groups were compared by Student’s t test, and the mean between multiple groups were compared by one-way ANOVA with Tukey’s correction for multiple hypothesis testing. A p-value less than 0.05 was considered to be a significant difference. All data were obtained from at least three independent experiments.

## Results

### OpA-induced cytotoxicity is selective to cancer cells and is potent *in vivo*

OpA has been demonstrated to possess cytotoxic activity towards various types of cancer cell lines, including human melanoma and glioblastoma (19, 21). In our previous work (22), we determined that epithelial-mesenchymal transition (EMT) enhances sensitivity to OpA and that breast cancer cell lines also exhibit differential sensitivity. We next sought to elucidate the molecular pathways involved in OpA-induced cytotoxicity across multiple breast cancer cell lines. We investigated the cytotoxicity of OpA in a panel of cell lines which showed that OpA exhibits significant cytotoxicity in breast cancer cell lines but lacks activity towards non-tumorigenic human cells (Fig.1A). The results obtained from the analysis suggest that OpA is capable of selectively targeting cancer cells without displaying any toxic behavior towards non-tumorigenic human breast cells (MCF10A) and human embryonic fibroblasts (293T). Corroborating these *in vitro* results, OpA, when dosed at 5 mg/kg, significantly reduces the tumor volume of MDA-MB-231 cells xenografted to immunocompromised mice, to a degree similar to docetaxel (Fig.1B). These findings suggest that OpA has the potential to serve as a treatment option for breast cancer patients; however, clarity as to the mechanism of action employed by OpA is needed.

**Figure 1.**
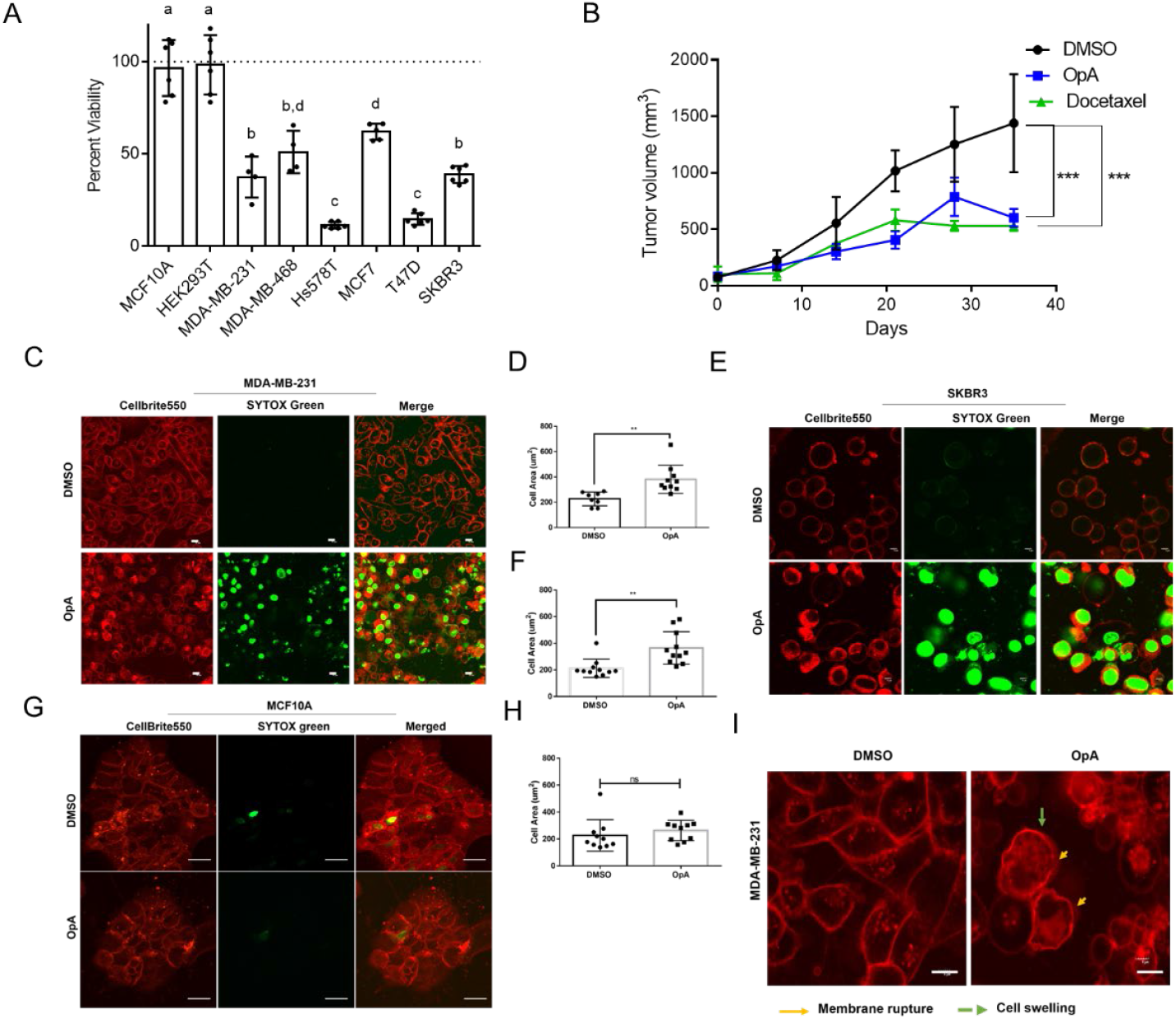
Ophiobolin A displays cancer cell-selective cytotoxicity via a lytic mechanism. (A) Cell viability of indicated cell lines treated with OpA at 500 nM for 24h. The values presented are relative to the viability of vehicle-treated cells and normalized to 100%. Letters indicate statistically significant groupings via one-way ANOVA with Tukey’s correction for multiple hypothesis testing. (B) MDA-MB-231 cells were injected into the mammary fat pad of Scid/bg mice. Upon reaching 100 mm^3^, mice were treated with 5 mg/kg OpA or 5 mg/kg docetaxel i.p. weekly. Tumor growth curve assessed by caliper measurement. Tumor size data compared by two-way ANOVA with Sidak’s correction for multiple hypothesis testing. (C-H) Images and quantifications showing plasma membrane stain (Cellbrite550) and DNA stain (SYTOX green) in OpA (500 nM) treated versus control cells for 6h in indicated cell lines. Original magnification 20X; scale bar 10 µm. Cell size measured as area using ImageJ software and compared by Student’s t-test. (I) Confocal images of MDA-MB-231 cells treated with OpA (500 nM) showing membrane rupture and cell swelling compared to DMSO control. Original magnification 20X; scale bar 5 µm. All data are presented as the mean ± SD from at least three independent experiments. *p< 0.05, **p<0.01, ***p<0.001, ****p<0.0001, ns=not significant vs control.

We next evaluated the mechanism by which OpA induces cytotoxicity in breast cancer cell lines. To compare the cytotoxic effect of OpA with a known cytotoxic drug, MBA-MB-231 cells were treated with OpA alone or in combination with docetaxel. Our results demonstrate that a combination of OpA and docetaxel displays a greater cytotoxic effect than each agent alone, indicating a potential complementary mode of action (SFig. 1A). In glioblastoma and melanoma, OpA has been shown to induce paraptosis and autophagy (19, 21). To test for paraptosis induction, we examined the capacity of spautin-1, an inhibitor of USP10 which is required for paraptosis (30), to rescue OpA-induced cytotoxicity. Our findings reveal that spautin-1 was ineffective in rescuing OpA-induced cytotoxicity against MDA-MB-231 cells and MCF7 cells (SFig. 1B), indicating a lack of dependency on USP10. We also assayed for swelling of the endoplasmic reticulum, an indication of paraptosis, by transmission electron microscopy (TEM). MDA-MB-231 cells failed to exhibit a significant increase in endoplasmic reticulum size (SFig. 1C/D). Similarly, we tested for evidence of the induction of autophagy through the detection of autophagolysosomes. While rapamycin was sufficient to cause an increase in autophagolysosome staining in MDA-MB-231 cells, OpA did not, (SFig. 1E) supporting the idea that autophagy is not induced.

We next evaluated the role of the caspase-dependent apoptotic cell death pathway in OpA-induced cytotoxicity by employing a pan-caspase inhibitor, zVAD. zVAD partially rescues caspase-dependent cell death in one of two ER-positive cell lines (T47D but not MCF7) and two of three TNBC cell lines (MDA-MB-231 and MDA-MB-468, but not Hs578T) (SFig. 2A), indicating that OpA-induced cell death involves caspase activation. However, the magnitude of rescue by zVAD was minimal, suggesting that OpA may induce cell death through additional pathways. Supporting this interpretation, MDA-MB-231 cells treated with OpA failed to show Annexin V-positivity, evident in staurosporine-treated cells (SFig. 2B/C). Additionally, OpA-treated MDA-MB-231 cells show no signs of membrane blebbing or apoptotic bodies as observed in the staurosporine-treated cells (SFig.2D).

### Ophiobolin A triggers morphological changes consistent with lytic cell death

To obtain morphological evidence of the cell death mechanism employed by OpA, we analyzed various breast cancer subtypes for structural changes in the plasma and nuclear membranes upon exposure to OpA by visualizing the plasma membrane and DNA. Both MDA-MB-231 (Fig. 1C) and SKBR3 (Fig. 1E) cell lines exhibit ruptured membranes and take up Sytox green, thus suggesting compromised membrane integrity consequent to OpA treatment. To assess the effect of OpA on non-transformed cells, MCF10A cells were treated with OpA and stained similarly. The results show that MCF10A cells are not affected by OpA treatment, as indicated by the intact membrane integrity (Fig. 1G). Moreover, cells that respond to OpA treatment exhibit a noticeably swollen appearance, as evidenced by quantification of cell area of MDA-MB-231 (Fig. 1D) and SKBR3 cells (Fig. 1F), but not MCF10A cells (Fig. 1H). This suggests that the cell-targeting ability of OpA is selective to cancer cells and does not affect non-cancer cells. Rupture of the plasma membrane is characteristic of induction of lytic cell death (31). Studies have documented the occurrence of plasma membrane rupture and cell swelling during both necroptosis and pyroptosis, two forms of cell death (31). Thus, we next sought to measure the capacity of OpA to activate features of these pathways.

### Ophiobolin A requires RIPK1 activity and can induce necrosome formation upon RIPK3 overexpression

Upon observing signs of lytic cell death, we opted to investigate necroptotic cell death (programmed necrosis), which acts through receptor-interacting serine/threonine-protein kinase 1 (RIPK1). We tested whether Necrostatin-1 (Nec-1), a RIPK1 inhibitor, was sufficient to rescue OpA-induced cytotoxicity. Our results demonstrate that Nec-1 blocks cell death in all the cell lines we evaluated, except for MCF7, which is less sensitive to OpA (Fig. 2A).

**Figure 2.**
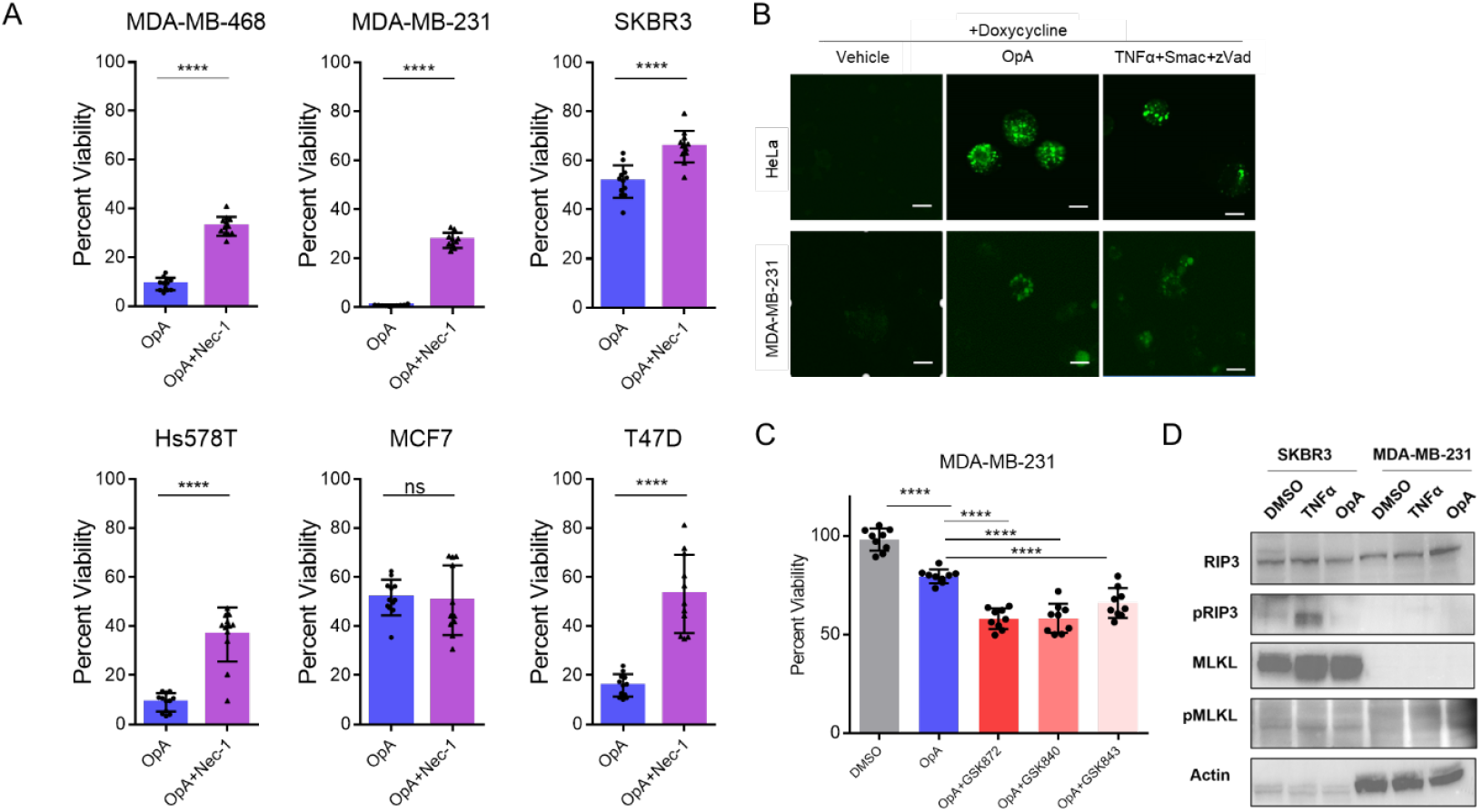
Ophiobolin A activity is dependent on RIPK1, but not RIPK3. (A) Viability of cells treated with 500 nM OpA in presence or absence of necrostatin-1 (100 µM). The values presented are relative to the viability of vehicle-treated cells and normalized to 100%. OpA+Nec-1 compared to OpA by Student’s t-test. (B) Confocal images of transiently transfected RIPK3-GFP MDA-MB-231 cells and stably transfected RIPK3-GFP HeLa cells treated with doxycycline (20 μg/ml) for 24h and with 500 nM OpA or 20 ng/ml TNFα or vehicle for 24h for visualization of the necrosome. Original magnification 20X; scale bar 20 µm (C) Viability of OpA-treated cells in presence or absence of RIPK3 inhibitors (GSK872, GSK840, GSK843) for 24h. The values presented are relative to the viability of DMSO-treated cells and normalized to 100%. All groups compared via one-way ANOVA with Tukey’s correction for multiple hypothesis testing. (D) Immunoblot analysis of total and phospho-RIPK3, total and phospho-MLKL using lysates of OpA- or TNFα-treated cells. β-Actin was used as an internal control. Cells were treated for 24h. Viability data are presented as the mean ± SD from at least three independent experiments. *p< 0.05, **p<0.01, ***p<0.001, ****p<0.0001, ns=not significant.

RIPK1 phosphorylates downstream kinase receptor-interacting serine/threonine-protein kinase 3 (RIPK3) which then interacts in a protein complex with mixed lineage kinase domain like pseudokinase (MLKL) in a structure termed the necrosome, which is a crucial step in inducing necroptotic cell death (32). The RIPK3 mRNA is typically expressed at lower levels in breast cancer cells when compared to normal cells, as per TCGA database, and can be silenced by DNA methylation (33). To examine the potential for OpA to induce necrosomes, we overexpressed GFP-tagged RIPK3 in MDA-MB-231 cells. The formation of puncta indicative of necrosomes was visualized via fluorescence microscopy. In contrast to vehicle control, OpA caused both MDA-MB-231 cells and HeLa cells expressing RIPK3-GFP to exhibit fluorescent puncta, similar to the formation of necrosomes induced by TNFα (13, 32) (Fig. 2B). HeLa cells expressing RIPK3-GFP were used in this study as a positive control, as the same plasmid used in this study and puncta (necrosome) shown in Sun et al., 2012 (32). However, MCF-7 cells overexpressing RIPK3 fail to exhibit puncta upon OpA treatment (SFig. 3A). Additionally, OpA induces intra-cellular calcium accumulation, a potential trigger of necroptosis as noted in other studies (34) (SFig. 3B). These data indicate that OpA relies upon RIPK1 for cytotoxic activity and has the capacity to induce necrosome formation in cancer cells upon overexpression of RIPK3.

To provide additional insight on the role of RIPK3 for OpA-induced cytotoxicity, we tested the effect of RIPK3 inhibitors GSK840, GSK843, and GSK870. However, all three inhibitors were unable to rescue cell death (Fig. 2C), indicating that endogenous RIPK3 may not be active in inducing necroptosis in response to OpA. To ascertain whether the kinase cascade characteristic of necroptosis is activated by OpA, we analyzed the phosphorylation of endogenous RIPK3 and MLKL proteins, using western blotting. Our data indicate that OpA was insufficient to induce the phosphorylation of RIPK3 and MLKL proteins in either MDA-MB-231 or SKBR3 cells (Fig. 2D), despite the capacity of TNFα to phosphorylate RIKP3 in SKBR3 cells. These data highlight the importance of RIPK1 activity in OpA-induced cytotoxicity yet are not supportive of canonical necroptosis induction.

### Ophiobolin A enhances the expression of cytokines and trigger inflammatory genes

In addition to necroptosis, lytic cell death may be accomplished through pyroptosis, which involves up-regulation and release of pro-inflammatory cytokines. Gene ontology (GO) analysis changes in mRNA expression upon OpA treatment of MDA-MB-231 cells shows elevated pathways including genes involved in cell cycle regulation, metabolism, as well as the response to interleukin-1 (Fig. 3A/B). Indeed, 28 of 34 genes in this category are elevated when compared to control (Fig. 3C). Additionally, NR4A1-NR4A3 (Nur77, Nurr1, and Nor-1), which are immediate early genes induced by growth factor, cytokines, inflammatory, physiological stimuli, and cellular stress (35), are highly activated in OpA-treated MDA-MB-231 cells (Fig. 3D) suggesting that OpA might activate pyroptosis.

**Figure 3.**
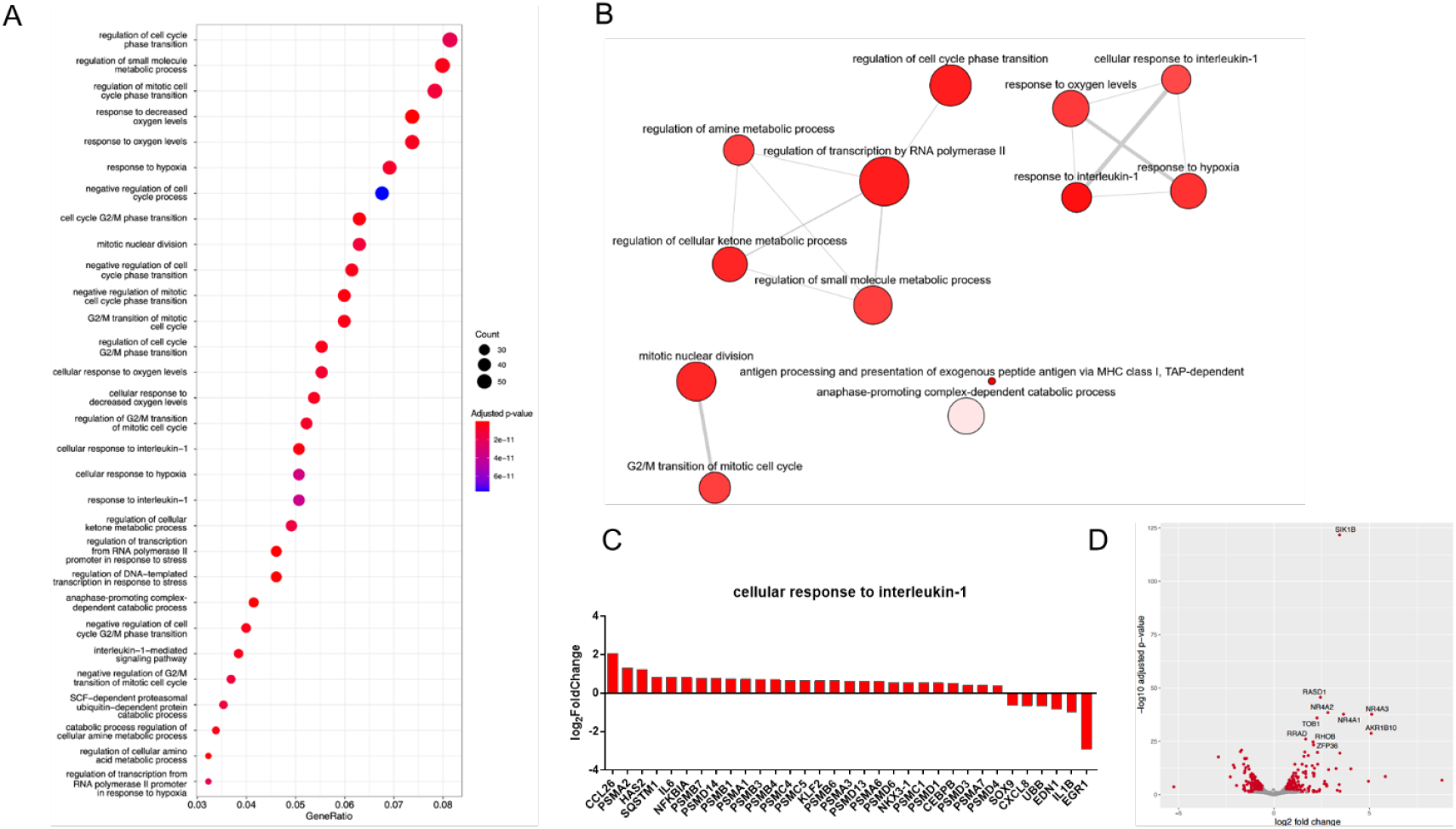
Ophiobolin A induces a gene expression signature indicative of response to cytokines. (A) Gene ontology (GO) analysis of differentially expressed genes from MDA-MB-231 cells treated with 250 nM OpA for 3h. (B) GO categories with p-values <10^−10^ were analyzed via Revigo to highlight non-redundant pathways. (C) Expression of select genes from RNA-seq data. (D) Volcano plot showing differently expressed genes from OpA-treated MDA-MB-231 cells.

To further test this hypothesis, we examined the ability of disulfiram, an inhibitor of Gasdermin D (GSDMD) cleavage and pyroptotic cell death (36, 37), to rescue OpA-induced cytotoxicity. Our data demonstrate that disulfiram exhibits a notable capacity to mitigate the cytotoxic effects induced by OpA in two of the three TNBC cell lines examined, although this effect was not observed in all cell lines (Fig. 4A). As upregulation of cytokine expression and secretion are key features of pyroptosis (38), we assessed the levels of three key cytokines: interleukin-6 (*IL6*), interleukin-8 (*CXCL8*), and interleukin-1 beta (*IL1B*). Our results reveal that OpA significantly elevates mRNA levels of inflammatory cytokines *CXCL8* in MDA-MB-468 cells, *IL6* and *IL1B* in MDA-MB-231 cells (Fig. 4C), and *IL6* and *CXCL8* in SKBR3 cells (Fig.4D), suggesting induction of pyroptosis by OpA.

**Figure 4.**
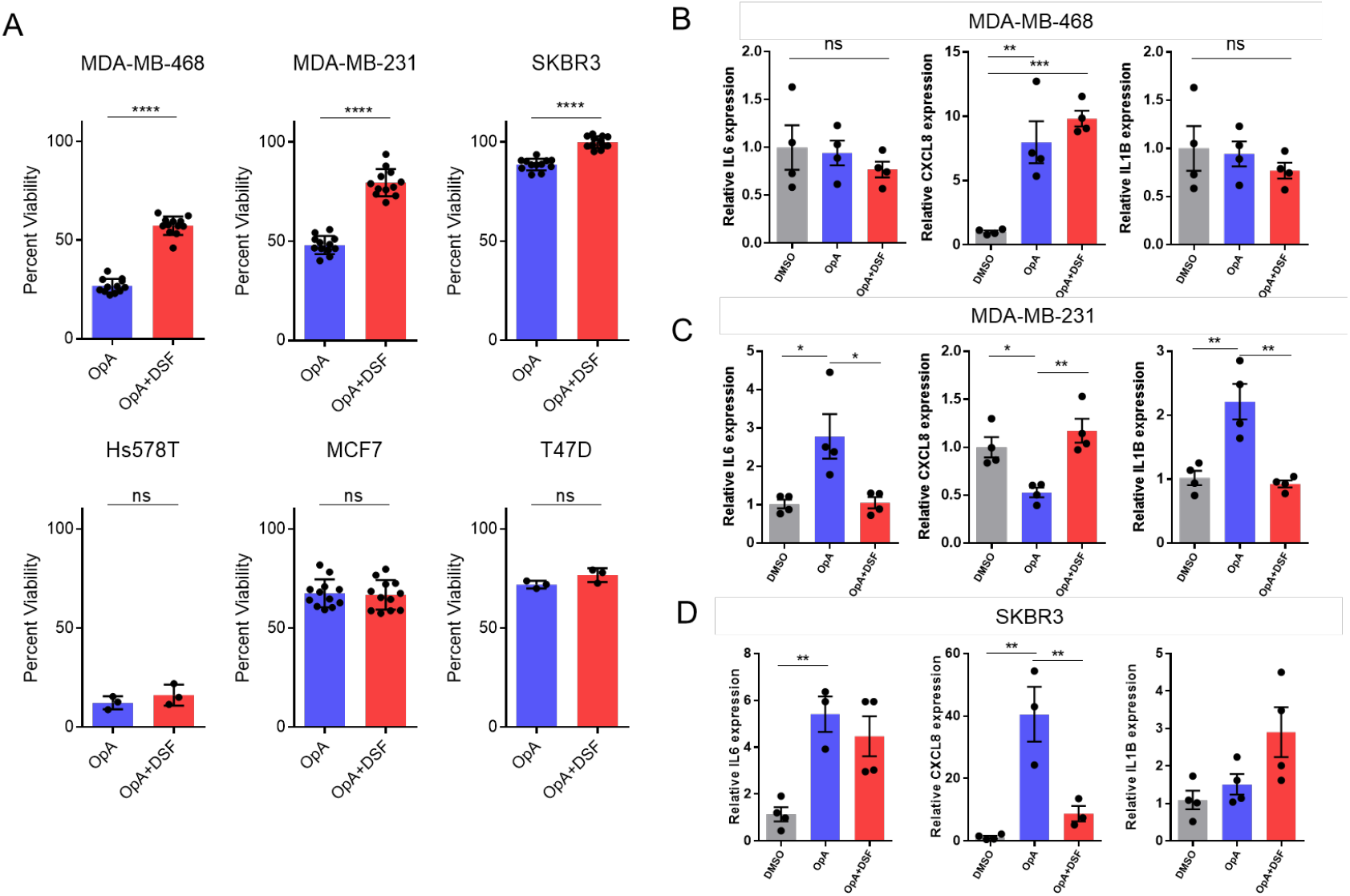
Ophiobolin A is blocked by disulfiram and enhances the expression of cytokines. (A) Viability of OpA-treated cells in presence or absence of disulfiram (1.25 µM) for 24h. (B-D) Real time qRT-PCR showing expression of indicated cytokines in OpA-treated cells with or without disulfiram for 6h for indicated cell lines. All data are presented as the mean ± SEM from at least three independent experiments using one way ANOVA with Tukey’s correction for multiple hypothesis testing. *p< 0.05, **p<0.01, ***p<0.001, ****p<0.0001, ns=not significant.

### Ophiobolin A facilitates release of cytokines and cleavage of Gasdermin D favoring pyroptotic cell death

Release of pro-inflammatory cytokines IL-8 and IL-1β has been attributed to a mechanism of pore formation in plasma membrane dependent on GSDMD (39). Cleavage of GSDMD and caspase-3 are well-established markers of pyroptotic cell death and necessary for membrane pore formation (40-42). Our western blot analysis reveals evidence of caspase-3 cleavage and GSDMD cleavage in SKBR3 cells (Fig. 5A), suggesting that OpA is capable of inducing pyroptotic cell death in HER2-positive cells. However, TNBC cell lines demonstrate neither caspase-3 nor GSDMD cleavage upon OpA treatment (Fig. 5A). To test for release of mature cytokines, we measured the protein concentration in conditioned media of cells treated with OpA. OpA significantly induces IL-8 release when compared to control in SKBR3 and MDA-MB-468 cells (Fig. 5B). Unexpectedly, disulfiram failed to reduce cytokine release. Altogether, our data suggest that, in SKBR3 cells, OpA induces transcription of pro-inflammatory cytokines, caspase-3 and GSDMD cleavage and cytokine release, while in TNBC cells, cytokine upregulation and release can occur in the absence of GSDMD cleavage. However, in all cell lines, cell death was dependent on the activity of RIPK1, indicative of PANoptosis (Fig. 5C).

**Figure 5.**
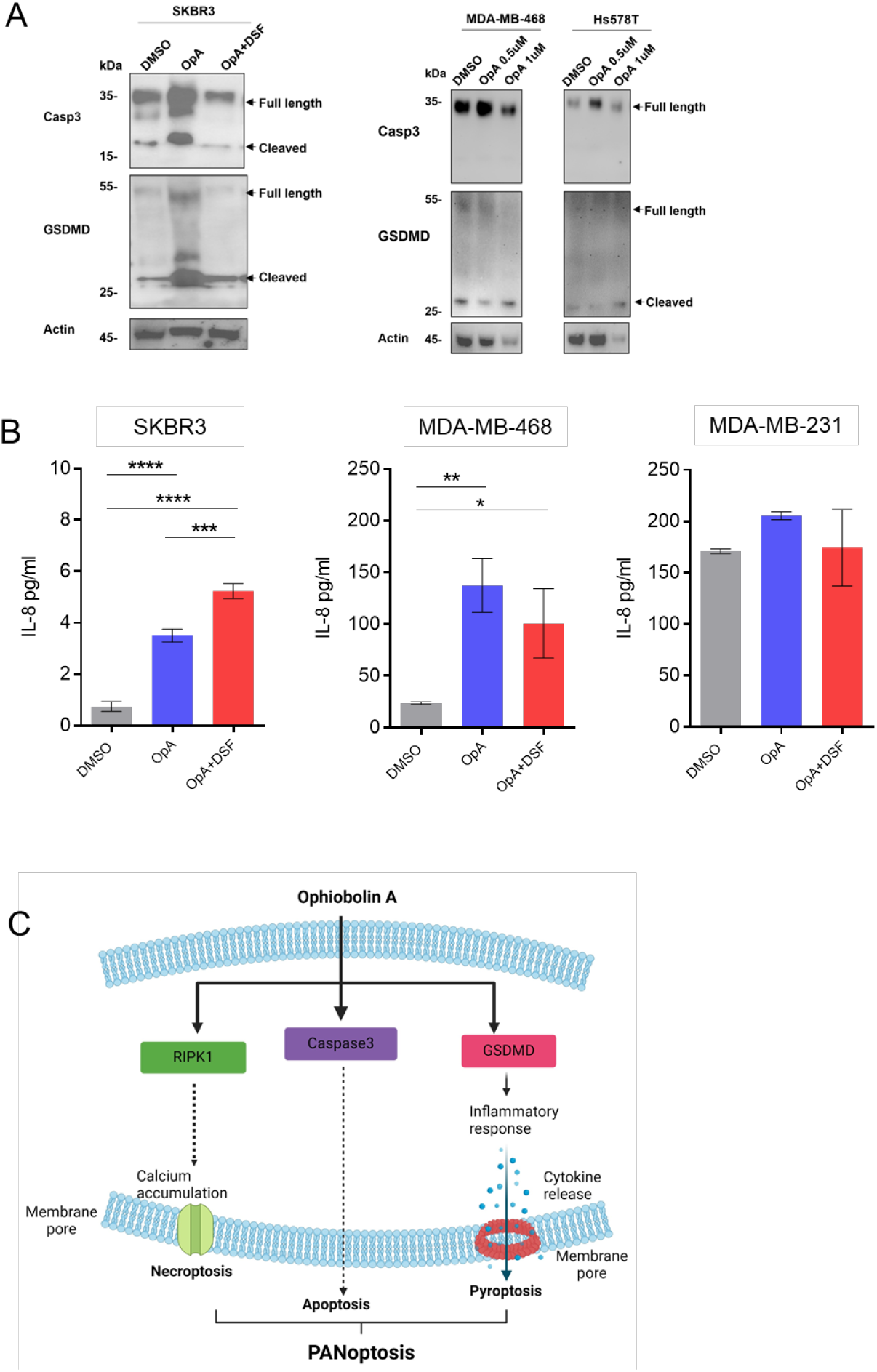
Ophiobolin A facilitates Gasdermin D cleavage and the release of cytokines. (A) Immunoblots showing the expression or cleavage of caspase 3 and GSDMD upon treatment with 500 nM OpA in the presence or absence of disulfiram for 12h (B) Levels of IL-8 protein released in conditioned media of indicated cell lines treated with OpA with or without disulfiram as in (A) was measured using ELISA. All data are presented as the mean ± SD from at least three independent experiments using one way ANOVA with Tukey’s correction for multiple hypothesis testing. (C) Graphical representation of the proposed mechanism by which OpA induces cell death in breast cancer cells. *p< 0.05, **p<0.01, ***p<0.001, ****p<0.0001, ns=not significant. The graphical illustration created using Biorender.com.

## Discussion

The ability of tumor cells to evade apoptosis is a critical factor in the development of cancer (43). Despite advancements in the management of breast cancer, the prognosis for patients with advanced disease remains unfavorable, mainly due to multidrug resistance (MDR) to cytotoxic chemotherapy drugs (2, 3). Therefore, identifying and characterizing additional therapeutic agents is critical in the fight against breast cancer.

Induction of non-apoptotic programmed cell death, including pyroptosis or PANoptosis, is likely to be essential for improvements in the treatment of various cancers. Our research has demonstrated that OpA triggers the formation of membrane pores and the uptake of Sytox Green, indicating the induction of lytic cell death in breast cancer cells, including TNBC. While OpA has been previously shown to induce paraptosis and autophagy in glioblastoma and melanoma cells (19, 21), our study did not reveal significant induction of paraptosis or autophagy in breast cancer cells. However, we did find that zVAD, a pan-caspase inhibitor rescues OpA-induced cell death in breast cancer cells despite the lack of annexin-V positivity. Therefore, our results suggest that OpA-induced cell death is mediated by caspase-dependent mechanisms in breast cancer cells but does not necessarily involve the apoptotic pathway.

Necroptosis and pyroptosis are two forms of programmed cell death that belong to the lytic pathway, characterized by the rupture of the cell membrane (44). Necroptosis, as its name suggests, is a programed form of necrosis, that functions through the activation of specific signaling pathways (45). RIPK1 is thought to be a crucial checkpoint regulator of apoptosis and necroptosis (46). RIPK1/RIPK3 necrosomes phosphorylate MLKL (mixed-lineage kinase domain-like protein), leading to the formation of membrane pores that promote necroptosis in the absence of caspase activation (47, 48). In our study, a RIPK1 inhibitor significantly blocked OpA-induced cell death in breast cancer cells, including TNBC, indicating that RIPK1 is involved in triggering the cell death pathway. Considering that RIPK3 is suppressed in tumor cells compared to normal cells, our RIPK3-overexpressing cells showed distinct puncta (necrosome) formation on OpA treatment, similar to the puncta shown by Sun et al. (32). However, OpA-treated cells failed to show phosphorylation of RIPK3 or MLKL in breast cancer cells. This indicates that OpA is able to initiate necroptosis in the abundance of endogenous RIPK3. In other words, a deficit of endogenous RIPK3 hinders OpA from triggering necroptosis.

While both pyroptosis and necroptosis are lytic cell death pathways with inflammatory characteristics, such as the release of damage-associated molecular patterns (DAMPs), they differ in their activation sequence. Pyroptosis can be a response to the presence of pathogen-associated molecular patterns (PAMPs) or DAMPs triggered by caspase-1 activation, resulting in the activation of cytokines IL-1β and IL-18 (6). In contrast, necroptosis is a secondary defense mechanism triggered by RIPK3 activation in response to the inhibition of pro-apoptotic caspase-8 (7). The two pathways are related in that DAMPs released during necroptosis can activate pyroptosis through regulation of the inflammatory axis, most notably IL-1α (5, 8, 9). Pyroptosis has been shown to be mediated by the cleavage of multiple gasdermin family members and the expression and release of inflammatory caspases IL-1β, IL-4, IL-5, and IL-11 (49, 50). Previous research has demonstrated that cleavage of casp3 results in the release of the N-terminal domain of GSDME, leading to the formation of pores in the cell membrane, cell swelling, rupture, and death (12). Furthermore, other studies have shown that cleavage of casp3 also induces pyroptosis through activation of GSDMD (51). In our study, we observed the activation of casp3 and cleavage of GSDMD, indicating that these two events may be responsible for the formation of membrane pores and cell swelling. The inhibition of caspases significantly blocked OpA-induced cell death, supporting this observation. Additionally, inhibition of GSDMD cleavage by disulfiram (36) (Fig. 5C) suppressed casp3 cleavage, indicating that both events are critical for the induction of pyroptosis.

OpA treatment also resulted in elevated transcription of cytokines IL-6, IL-8, and IL-1β and a significant release of IL-8, as observed in studies of pyroptosis (39). Furthermore, GO analysis of RNA-seq data revealed an elevated interleukin-1-mediated signaling pathway. Additionally, differential expression of NR4A1-NR4A3 genes was observed in OpA-treated TNBC cells, suggesting that OpA induces inflammation (35). The results of this study demonstrate that OpA treatment leads to the differential expression of NR4A1-NR4A3 genes, indicating a pro-inflammatory response in TNBC cells.

Our study reveals that OpA induces typical pyroptotic characteristics, including cell membrane rupture, cell swelling, the release of cytokines, and GSDMD cleavage in breast cancer cells. However, our research also raises the question of how RIPK1 is involved in OpA-induced pyroptosis. In line with previous observations (52), we speculate that there could be a crosstalk between necroptosis and pyroptosis in inducing cell death in breast cancer cells. Such crosstalk between cell death pathways has been recently reported in other studies, where apoptosis, necroptosis, and pyroptosis interact to activate inflammatory cell death, in a process termed PANoptosis (48, 52-55). In PANoptosis, despite RIPK3-MLKL hyperactivation, RIPK1 efficiently drives inflammasome activation and IL-18 secretion (52). The activation of casp3, RIPK1, and GSDMD suggests that PANoptosis could be activated upon OpA treatment in breast cancer.

In summary, our study uncovers a novel mechanism by which OpA induces PANoptosis in breast cancer cells via RIPK1 and caspase-3-mediated GSDMD cleavage. We found that OpA triggers pyroptosis-like cell death in breast cancer cells, characterized by cellular swelling, membrane rupture, and cytokine release, and is capable of inducing necrosomes in RIPK3-abundant breast cancer cells. These findings indicate that OpA may be a promising therapeutic candidate for the treatment of breast cancer. Overall, our results imply that OpA treatment may have therapeutic potential for TNBC by inducing PANoptotic cell death and potentially synergize with immunotherapy.

## AUTHOR CONTRIBUTIONS

SR and JHT designed the study. SR, TO, AS, JT, and KHL performed experiments. AK, AB, AE, and DR provided key reagents. MLB analyzed RNA-seq data. DR, AK, and JHT obtained funding. SR and JHT confirm the authenticity of all the raw data. All authors read and approved the final manuscript.

## PATIENT CONSENT FOR PUBLICATION

Not applicable

## DATA SHARING

RNA-seq data is publicly available by the Gene Expression Omnibus hosting by NIH at accession GSE277026. Other data will be made available upon reasonable request.

## ETHICS APPROVAL

Project carried out under the approval of Baylor University IACUC according to protocol 1441130.

## ACKNOWLEDGEMENTS

The authors thank Dr. Zhigao Wang, Associate professor, Center for Regenerative medicine, University of South Florida, for kindly providing RIPK3-GFP plasmids. SR thanks Dr. Bernd Zechmann, Director, Center for Microscopy and Imaging and Dr. Dwayne Simmons, Professor, Department of Biology, Baylor University for support with TEM and confocal imaging.

## REFERENCES

1. Sung H, Ferlay J, Siegel RL, Laversanne M, Soerjomataram I, Jemal A, et al. Global Cancer Statistics 2020: GLOBOCAN Estimates of Incidence and Mortality Worldwide for 36 Cancers in 185 Countries. CA Cancer J Clin. 2021;71(3):209–49.

2. Lim B, Greer Y, Lipkowitz S, Takebe N. Novel Apoptosis-Inducing Agents for the Treatment of Cancer, a New Arsenal in the Toolbox. Cancers (Basel). 2019;11(8).

3. Reed JC. Apoptosis-based therapies. Nat Rev Drug Discov. 2002;1(2):111–21.

4. O’Reilly EA, Gubbins L, Sharma S, Tully R, Guang MH, Weiner-Gorzel K, et al. The fate of chemoresistance in triple negative breast cancer (TNBC). BBA Clin. 2015;3:257–75.

5. Ouyang L, Shi Z, Zhao S, Wang FT, Zhou TT, Liu B, et al. Programmed cell death pathways in cancer: a review of apoptosis, autophagy and programmed necrosis. Cell Prolif. 2012;45(6):487–98.

6. Galluzzi L, Vitale I, Aaronson SA, Abrams JM, Adam D, Agostinis P, et al. Molecular mechanisms of cell death: recommendations of the Nomenclature Committee on Cell Death 2018. Cell Death Differ. 2018;25(3):486–541.

7. Vitale I, Pietrocola F, Guilbaud E, Aaronson SA, Abrams JM, Adam D, et al. Apoptotic cell death in disease-Current understanding of the NCCD 2023. Cell Death Differ. 2023;30(5):1097–154.

8. Yu Y, Yan Y, Niu F, Wang Y, Chen X, Su G, et al. Ferroptosis: a cell death connecting oxidative stress, inflammation and cardiovascular diseases. Cell Death Discov. 2021;7(1):193.

9. Hanson S, Dharan A P VJ, Pal S, Nair BG, Kar R, et al. Paraptosis: a unique cell death mode for targeting cancer. Front Pharmacol. 2023;14:1159409.

10. Chen C, Ye Q, Wang L, Zhou J, Xiang A, Lin X, et al. Targeting pyroptosis in breast cancer: biological functions and therapeutic potentials on It. Cell Death Discov. 2023;9(1):75.

11. Jorgensen I, Miao EA. Pyroptotic cell death defends against intracellular pathogens. Immunol Rev. 2015;265(1):130–42.

12. al JMe. The casp-3/GSDME signal pathway as a switch between apoptosis and pyroptosis.

13. Wang Y, Gao W, Shi X, Ding J, Liu W, He H, et al. Chemotherapy drugs induce pyroptosis through caspase-3 cleavage of a gasdermin. Nature. 2017;547(7661):99–103.

14. Gao J, Xiong A, Liu J, Li X, Wang J, Zhang L, et al. PANoptosis: bridging apoptosis, pyroptosis, and necroptosis in cancer progression and treatment. Cancer Gene Ther. 2024;31(7):970–83.

15. Bladt TT, Durr C, Knudsen PB, Kildgaard S, Frisvad JC, Gotfredsen CH, et al. Bio-activity and dereplication-based discovery of ophiobolins and other fungal secondary metabolites targeting leukemia cells. Molecules. 2013;18(12):14629–50.

16. Tao Y, Reisenauer KN, Masi M, Evidente A, Taube JH, Romo D. Pharmacophore-Directed Retrosynthesis Applied to Ophiobolin A: Simplified Bicyclic Derivatives Displaying Anticancer Activity. Org Lett. 2020;22(21):8307–12.

17. Masi M, Dasari R, Evidente A, Mathieu V, Kornienko A. Chemistry and biology of ophiobolin A and its congeners. Bioorg Med Chem Lett. 2019;29(7):859–69.

18. Reisenauer KN, Aroujo J, Tao YF, Ranganathan S, Romo D, Taube JH. Therapeutic vulnerabilities of cancer stem cells and effects of natural products. Nat Prod Rep. 2023;40(8):1432–56.

19. Bury M, Girault A, Megalizzi V, Spiegl-Kreinecker S, Mathieu V, Berger W, et al. Ophiobolin A induces paraptosis-like cell death in human glioblastoma cells by decreasing BKCa channel activity. Cell Death Dis. 2013;4(3):e561.

20. Morrison R, Lodge T, Evidente A, Kiss R, Townley H. Ophiobolin A, a sesterpenoid fungal phytotoxin, displays different mechanisms of cell death in mammalian cells depending upon the cancer cell origin. Int J Oncol. 2017;50(3):773–86.

21. Rodolfo C, Rocco M, Cattaneo L, Tartaglia M, Sassi M, Aducci P, et al. Ophiobolin A Induces Autophagy and Activates the Mitochondrial Pathway of Apoptosis in Human Melanoma Cells. PLoS One. 2016;11(12):e0167672.

22. Reisenauer KN, Tao Y, Das P, Song S, Svatek H, Patel SD, et al. Epithelial-mesenchymal transition sensitizes breast cancer cells to cell death via the fungus-derived sesterterpenoid ophiobolin A. Sci Rep. 2021;11(1):10652.

23. Sun LM, Wang HY, Wang ZG, He SD, Chen S, Liao DH, et al. Mixed Lineage Kinase Domain-like Protein Mediates Necrosis Signaling Downstream of RIP3 Kinase. Cell. 2012;148(1-2):213–27.

24. Evidente A, Andolfi A, Cimmino A, Vurro M, Fracchiolla M, Charudattan R. Herbicidal potential of ophiobolins produced by Drechslera gigantea. J Agric Food Chem. 2006;54(5):1779–83.

25. Heid CA, Stevens J, Livak KJ, Williams PM. Real time quantitative PCR. Genome Res. 1996;6(10):986–94.

26. Patro R, Duggal G, Love MI, Irizarry RA, Kingsford C. Salmon provides fast and bias-aware quantification of transcript expression. Nature methods. 2017;14(4):417–9.

27. Love MI, Huber W, Anders S. Moderated estimation of fold change and dispersion for RNA-seq data with DESeq2. Genome Biol. 2014;15(12):550.

28. Love MI, Soneson C, Hickey PF, Johnson LK, Pierce NT, Shepherd L, et al. Tximeta: Reference sequence checksums for provenance identification in RNA-seq. PLoS Comput Biol. 2020;16(2):e1007664.

29. Wu T, Hu E, Xu S, Chen M, Guo P, Dai Z, et al. clusterProfiler 4.0: A universal enrichment tool for interpreting omics data. Innovation (Camb). 2021;2(3):100141.

30. Kim JY, Lee DM, Woo HG, Kim KD, Lee HJ, Kwon YJ, et al. RNAi Screening-based Identification of USP10 as a Novel Regulator of Paraptosis. Scientific reports. 2019;9(1):4909.

31. Zhang Y, Chen X, Gueydan C, Han J. Plasma membrane changes during programmed cell deaths. Cell Res. 2018;28(1):9–21.

32. Sun L, Wang H, Wang Z, He S, Chen S, Liao D, et al. Mixed lineage kinase domain-like protein mediates necrosis signaling downstream of RIP3 kinase. Cell. 2012;148(1-2):213–27.

33. Koo GB, Morgan MJ, Lee DG, Kim WJ, Yoon JH, Koo JS, et al. Methylation-dependent loss of RIP3 expression in cancer represses programmed necrosis in response to chemotherapeutics. Cell Res. 2015;25(6):707–25.

34. Nomura M, Ueno A, Saga K, Fukuzawa M, Kaneda Y. Accumulation of cytosolic calcium induces necroptotic cell death in human neuroblastoma. Cancer Res. 2014;74(4):1056–66.

35. Crean D, Murphy EP. Targeting NR4A Nuclear Receptors to Control Stromal Cell Inflammation, Metabolism, Angiogenesis, and Tumorigenesis. Front Cell Dev Biol. 2021;9:589770.

36. Hu JJ, Liu X, Xia S, Zhang Z, Zhang Y, Zhao J, et al. FDA-approved disulfiram inhibits pyroptosis by blocking gasdermin D pore formation. Nat Immunol. 2020;21(7):736–45.

37. Zhang Y, Zhang R, Han X. Disulfiram inhibits inflammation and fibrosis in a rat unilateral ureteral obstruction model by inhibiting gasdermin D cleavage and pyroptosis. Inflamm Res. 2021;70(5):543–52.

38. Li S, Bracey S, Liu Z, Xiao TS. Regulation of gasdermins in pyroptosis and cytokine release. Adv Immunol. 2023;158:75–106.

39. Chen M, Rong R, Xia X. Spotlight on pyroptosis: role in pathogenesis and therapeutic potential of ocular diseases. J Neuroinflammation. 2022;19(1):183.

40. Chen X, He WT, Hu L, Li J, Fang Y, Wang X, et al. Pyroptosis is driven by non-selective gasdermin-D pore and its morphology is different from MLKL channel-mediated necroptosis. Cell Res. 2016;26(9):1007–20.

41. Chu X, Xiao X, Wang G, Uosef A, Lou X, Arnold P, et al. Gasdermin D-mediated pyroptosis is regulated by AMPK-mediated phosphorylation in tumor cells. Cell Death Dis. 2023;14(7):469.

42. Devant P, Borsic E, Ngwa EM, Xiao H, Chouchani ET, Thiagarajah JR, et al. Gasdermin D pore-forming activity is redox-sensitive. Cell Rep. 2023;42(1):112008.

43. Igney FH, Krammer PH. Death and anti-death: tumour resistance to apoptosis. Nat Rev Cancer. 2002;2(4):277–88.

44. Lawrence SM, Corriden R, Nizet V. How Neutrophils Meet Their End. Trends Immunol. 2020;41(6):531–44.

45. Xia X, Lei L, Wang S, Hu J, Zhang G. Necroptosis and its role in infectious diseases. Apoptosis. 2020;25(3-4):169–78.

46. Annibaldi A, Meier P. Checkpoints in TNF-Induced Cell Death: Implications in Inflammation and Cancer. Trends Mol Med. 2018;24(1):49–65.

47. Li J, McQuade T, Siemer AB, Napetschnig J, Moriwaki K, Hsiao YS, et al. The RIP1/RIP3 necrosome forms a functional amyloid signaling complex required for programmed necrosis. Cell. 2012;150(2):339–50.

48. Gao J, Xiong A, Liu J, Li X, Wang J, Zhang L, et al. PANoptosis: bridging apoptosis, pyroptosis, and necroptosis in cancer progression and treatment. Cancer Gene Ther. 2024.

49. Ding J, Wang K, Liu W, She Y, Sun Q, Shi J, et al. Pore-forming activity and structural autoinhibition of the gasdermin family. Nature. 2016;535(7610):111–6.

50. Wei S, Feng M, Zhang S. Molecular Characteristics of Cell Pyroptosis and Its Inhibitors: A Review of Activation, Regulation, and Inhibitors. Int J Mol Sci. 2022;23(24).

51. Rogers C, Fernandes-Alnemri T, Mayes L, Alnemri D, Cingolani G, Alnemri ES. Cleavage of DFNA5 by caspase-3 during apoptosis mediates progression to secondary necrotic/pyroptotic cell death. Nat Commun. 2017;8:14128.

52. Malireddi RKS, Kesavardhana S, Karki R, Kancharana B, Burton AR, Kanneganti TD. RIPK1 Distinctly Regulates Yersinia-Induced Inflammatory Cell Death, PANoptosis. Immunohorizons. 2020;4(12):789–96.

53. Sun X, Yang Y, Meng X, Li J, Liu X, Liu H. PANoptosis: Mechanisms, biology, and role in disease. Immunol Rev. 2024;321(1):246–62.

54. Christgen S, Zheng M, Kesavardhana S, Karki R, Malireddi RKS, Banoth B, et al. Identification of the PANoptosome: A Molecular Platform Triggering Pyroptosis, Apoptosis, and Necroptosis (PANoptosis). Front Cell Infect Microbiol. 2020;10:237.

55. Lee S, Karki R, Wang Y, Nguyen LN, Kalathur RC, Kanneganti TD. AIM2 forms a complex with pyrin and ZBP1 to drive PANoptosis and host defence. Nature. 2021;597(7876):415–9.

