## Supplemental Figures for "PANoptosis, a combination of inflammatory cell death mechanisms, induced by Ophiobolin A in breast cancer cell lines"

^1^Department of Biology, Baylor University, Waco, TX, 76706, USA, ^2^Department of Chemistry and Biochemistry, Texas State University, San Marcos, TX, 78666, USA, ^3^Institute of Sciences and Food Production, National Research Council, 70126, Bari, Italy, ^4^Institute of Biomolecular Sciences, National Research Council, 80078, Pozzuoli, Italy, ^5^Department of Computer Science, Baylor University, Waco, TX, 76706, USA, ^6^Department of Chemistry and Biochemistry, Baylor University, Waco, TX, 76706 USA.

Running title: Ranganathan et al. OpA induces PANoptosis

Keywords: breast cancer, ophiobolin A, PANoptosis, pyroptosis, cell death

#Corresponding author

Dr. Joseph Taube, One Bear Place #97388, Waco, TX, 76706 USA

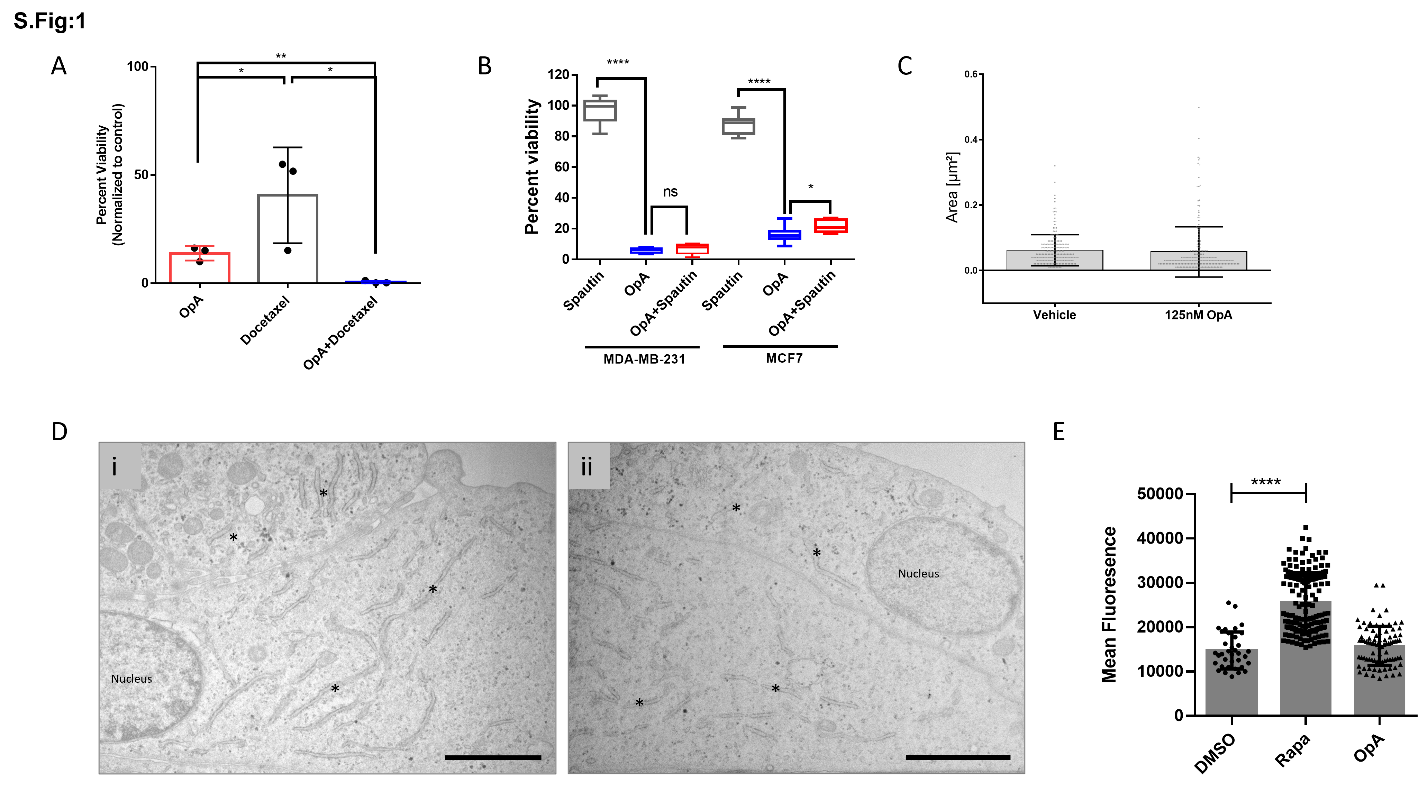

**SFig.1.** (A) Viability of MDA-MB-231 cells treated with OpA, docetaxel, or a combination for 24h. (B) Viability of OpA-treated cells in presence or absence of spautin-1 (1 µM) for 24h. The values presented are relative to the viability of untreated cells and normalized to 100%. (C/D) Quantification and images of TEM micrographs of MDA-MB 231 treated with vehicle (i) and 125 nM OpA (ii). Scale bar represents 2 µm, endoplasmic reticulum indicated by (*) (E) MDA-MB-231 cells were plated on glass-bottom slides for imaging, then treated with OpA or rapamycin for 24h before staining for autophagolysosomes using CytoID. Intensity of stain of individual cells in plotted. *p< 0.05, **p<0.01, ***p<0.001, ****p<0.0001, ns=not significant vs control using Student’s t-test.

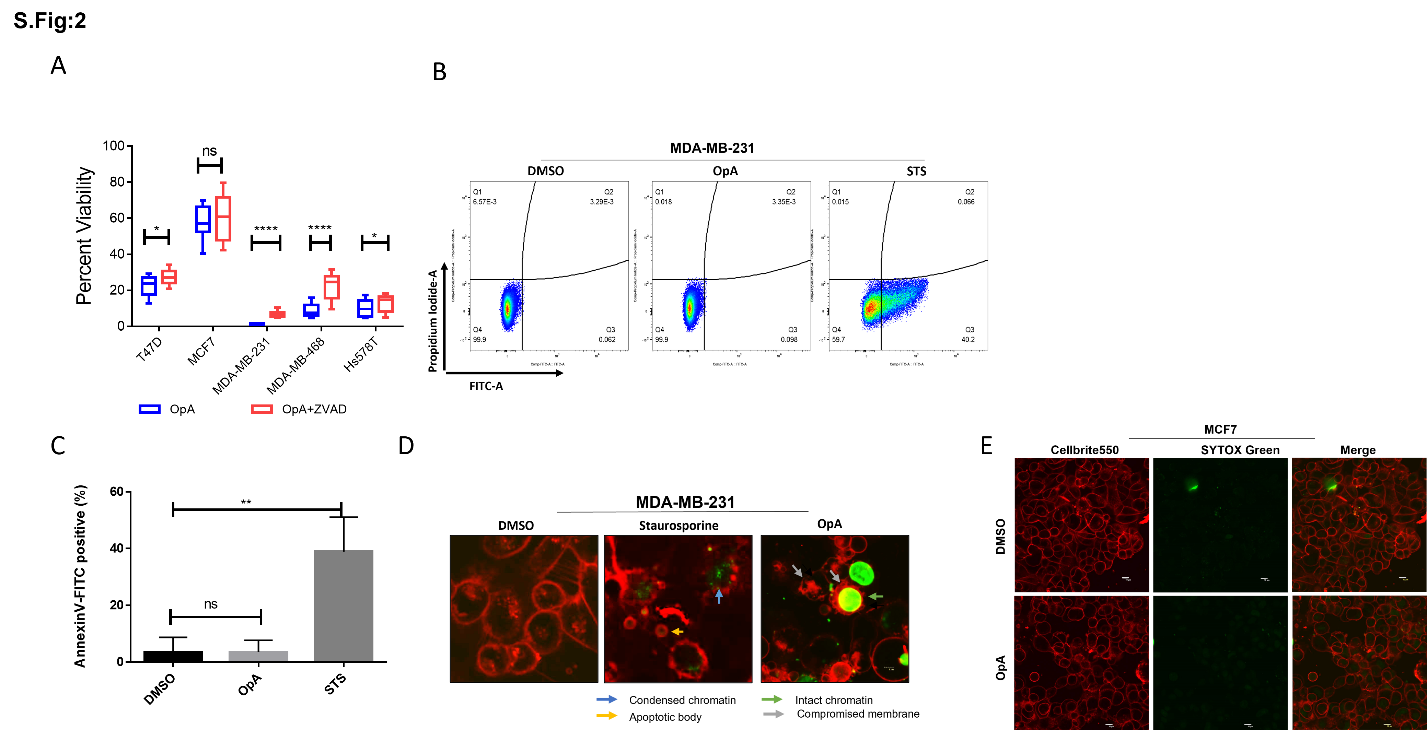
**SFig. 2.** (A) Cell viability of OpA-treated cells in presence or absence of zVAD for 24h. The values presented are relative to the viability of untreated cells and normalized to 100%. (B/C) Plot and quantification of flow cytometry analysis showing Annexin V-FITC positive cells in MDA-MB-231 cells treated with OpA or staurosporine (0.5 µM) (STS). (D) Confocal images showing apoptotic bodies and chromatin condensation in STS- but not OpA-treated MDA-MB-231 cells. (E) Confocal microscopy images of MCF7 cells showing plasma membrane stain (Cellbrite550) and lack of DNA stain (SYTOX green) in OpA (500 nM) treated versus control cells for 6h. Graphed data are presented as the mean ± SD from three independent experiments. *p< 0.05, **p<0.01, ***p<0.001, ****p<0.0001, ns=not significant vs control using Student’s t-test.

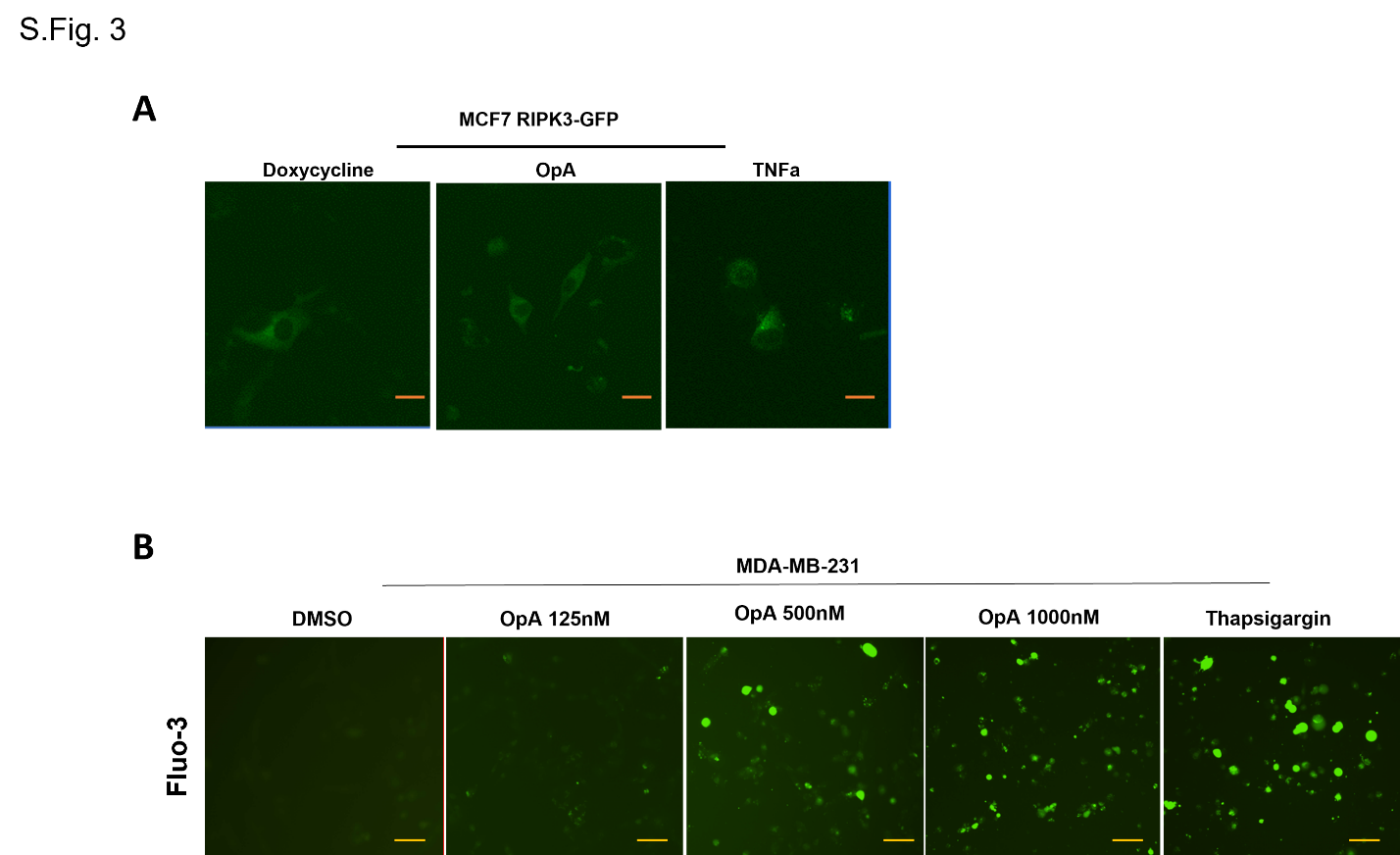

**SFig. 3.** (A) Confocal images of MCF7 RIPK3-GFP cells showing GFP expression but no puncta upon OpA treatment for 24h. (B) Representative epi-fluorescence images of MDA-MB-231 cells showing calcium accumulation upon increasing dosage of OpA for 24h.
